# Identification of Proliferation-Specific Dependencies for Therapeutic Targeting of Liver Cancer

**DOI:** 10.64898/2026.07.09.737474

**Authors:** Mirco Castoldi

## Abstract

Hepatocellular carcinoma (HCC) remains a leading cause of cancer-related mortality worldwide despite recent therapeutic advances, driven in part by its marked etiological and molecular heterogeneity and the lack of broadly effective therapeutic targets. Identifying conserved tumor dependencies shared across distinct etiological backgrounds may provide new opportunities for targeted therapy. Here, we developed an integrative computational framework to systematically integrate transcriptomic, functional genomics, and clinical datasets for the identification and prioritization of candidate tumor dependency genes in liver cancer. We reanalyzed transcriptomic data from murine models of liver cancer driven by genotoxic (DEN), oncogenic (c-Myc), and inflammatory (lymphotoxin) stimuli, identifying more than 380 genes consistently upregulated across all tumor models. Functional enrichment analysis revealed a strong overrepresentation of cell cycle-related pathways and liver cancer signatures. Integration with DepMap dependency datasets identified 26 genes with strong dependency scores. Candidate genes were further prioritized by comparing their expression across models of liver regeneration, chronic liver injury, and liver cancer. Analysis of the TCGA-LIHC cohort confirmed significant overexpression of all 26 genes in human HCC, with high expression associated with poor patient survival. Together, these findings establish an integrative framework for identifying conserved tumor dependencies, providing a prioritized set of proliferation-associated genes for functional evaluation as therapeutic targets in HCC.

## Introduction

Hepatocellular carcinoma (HCC) is the most common primary liver malignancy and the third leading cause of cancer-related death worldwide^1^. Despite considerable advances in surveillance, diagnosis, and systemic therapies, including multikinase inhibitors and immune checkpoint inhibitors, the prognosis for patients with advanced HCC remains poor^2^. HCC arises from diverse etiological factors, including chronic viral hepatitis^3^, alcohol-associated liver disease^4^, and metabolic dysfunction-associated steatotic liver disease^5^, which converge on common pathological outcomes such as chronic inflammation, fibrosis, and dysregulated proliferation of liver cells. This remarkable biological heterogeneity remains a major challenge to the development of broadly effective targeted therapies. An attractive strategy for overcoming this heterogeneity is the identification of conserved tumor dependency genes. Unlike oncogenic driver mutations, tumor dependencies represent genes that are required for tumor cell survival or proliferation regardless of the initiating genetic or environmental insult. Such genes may therefore constitute shared vulnerabilities across molecularly distinct tumors. Moreover, dependency genes that are preferentially active in proliferating cells could provide a therapeutic window for selectively targeting malignant hepatocytes while minimizing toxicity to the largely quiescent healthy liver.

The rapid expansion of publicly available transcriptomic, functional genomics, and clinical datasets has created unprecedented opportunities to systematically identify these conserved vulnerabilities. Resources such as transcriptomic repositories (GEO^6^, ArrayExpress^7^), genome-wide CRISPR and RNAi dependency screens (DepMap^8^), and large patient cohorts (TCGA) collectively capture complementary aspects of tumor biology. However, these datasets are frequently analyzed independently, making it difficult to distinguish context-specific molecular alterations from fundamental tumor dependencies. Integrative and reproducible analytical strategies capable of combining heterogeneous data across experimental systems and clinical cohorts are therefore needed to improve therapeutic target prioritization.

Here, we developed a computational framework based on custom R pipelines that systematically integrates transcriptomic, functional genomics, and clinical datasets to identify conserved tumor dependency genes in hepatocellular carcinoma. As a proof of concept, we applied this workflow to multiple murine models of hepatocarcinogenesis^9, 10^ representing distinct etiological mechanisms and prioritized genes that were consistently induced across tumor models and associated with supporting the fitness of proliferating cancer cells. We further incorporated proliferative state as a biological selection criterion by comparing candidate gene expression across models of physiological liver regeneration and chronic liver injury, thereby distinguishing transient regenerative responses from persistent tumor-associated activation. Finally, we validated the clinical relevance of prioritized candidates in human HCC cohorts through analyses of gene expression and patient outcome. Beyond identifying candidate therapeutic targets, our approach demonstrates how the integration of complementary public datasets can accelerate translational target discovery and provides a resource for future studies aimed at developing selective therapeutic strategies for liver cancer.

## Results

### Identification of conserved transcriptional alterations across etiologically distinct murine models of liver cancer

To identify genes consistently deregulated across distinct etiological models of liver cancer, we reanalyzed a dataset that was previously generated in our laboratory and that if is retrievable via Gene Expression Omnibus (GEO) deposited dataset GSE102416^10^. The dataset comprises three complementary murine models of liver cancer representing distinct pathogenic mechanisms: genotoxic (DEN), oncogenic (c-Myc overexpression), and inflammatory (lymphotoxin overexpression). For each model, RNA was isolated from normal liver (CTRL), non-tumorous liver adjacent to tumors (NTL), and tumor tissue. Transcriptomic data were processed using our custom easyGEOarray (eGA) pipeline, and differential expression analysis comparing tumor with CTRL and NTL samples identified more than 380 significantly upregulated and 180 significantly downregulated genes (Figure 1). As the objective of this study was to identify candidate tumor dependencies that may represent therapeutically targetable vulnerabilities, subsequent analyses focused exclusively on the upregulated gene set. Functional disease enrichment analysis using DisGeNET^11^ identified liver fibrosis as the most significantly enriched disease term, consistent with the central role of fibrosis in hepatocarcinogenesis and supporting the biological relevance of the identified transcriptional signature. Moreover, gene ontology (GO) analysis identified a significant enrichment for term associate to cell division, mitotic spindle assembly and motor proteins.

**Figure 1.**
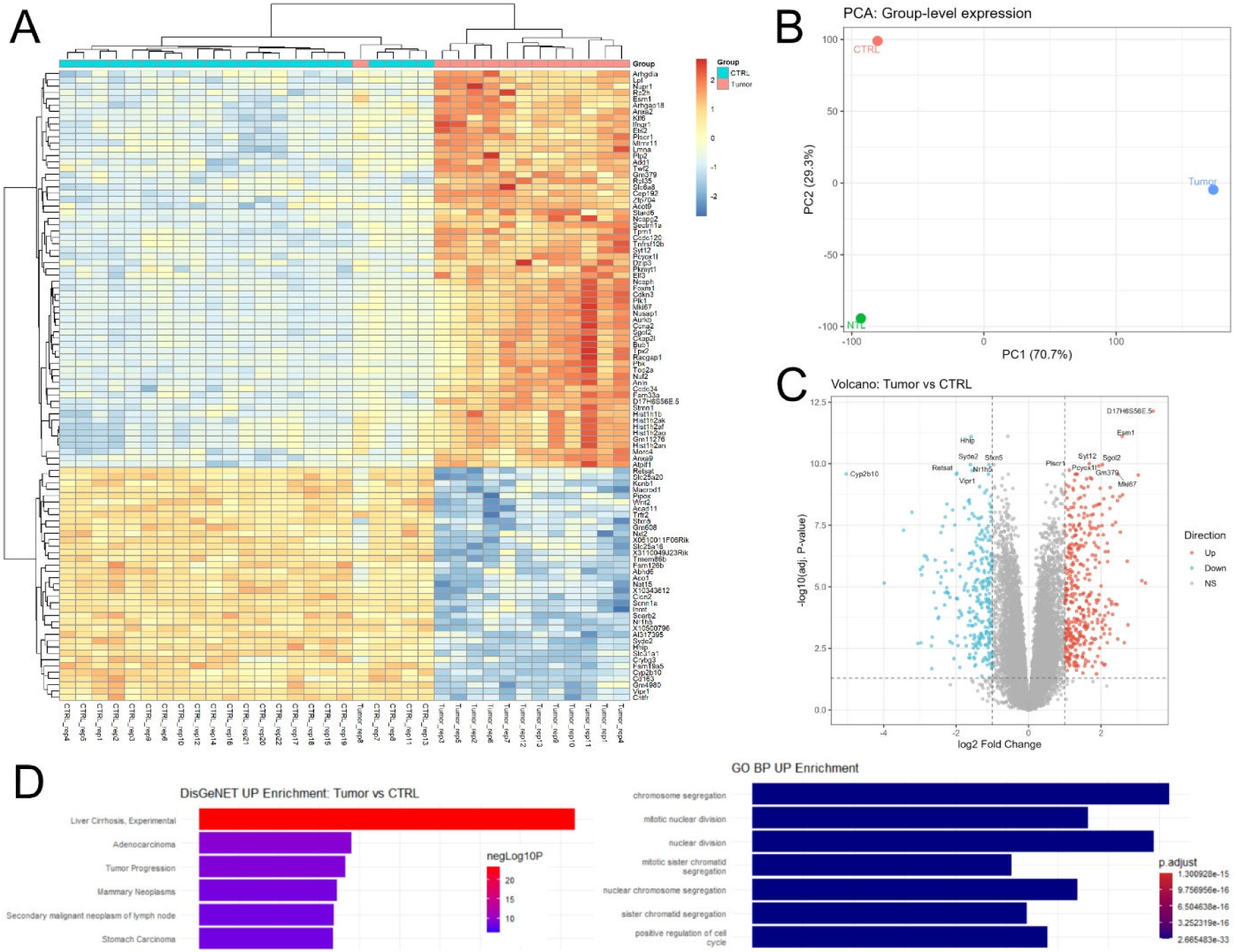
Identification of conserved transcriptional alterations across etiologically distinct murine HCC models. (**A**) Hierarchical clustering of differentially expressed genes across three complementary mouse models of liver cancer (GSE102416): genotoxic (DEN), oncogenic (c-Myc overexpression), and inflammatory (lymphotoxin overexpression). Tumor samples were compared with normal liver (CTRL) and non-tumorous liver adjacent to tumors (NTL) using the easyGEOarray (eGA) pipeline. (**B**) Principal component analysis (PCA) and (**C**) volcano plot showing significantly upregulated and downregulated genes. (**D**) Disease enrichment analysis of significantly upregulated genes using DisGeNET (left) identified “liver fibrosis” as the most significantly enriched disease term. Gene Ontology biological process analysis (GO-BP, right) showed significant enrichment of terms associated with cell division.

### Functional prioritization of conserved tumor-associated genes using DepMap

To prioritize candidate genes with functional relevance for tumor cell survival, the 380 conserved upregulated genes were interrogated using our custom easyDepMap (eDepMap) pipeline. The pipeline queries the Cancer Dependency Map (DepMap), a functional genomics resource developed by the Broad Institute (https://depmap.org) that integrates genome-wide CRISPR-Cas9 loss-of-function with RNAi-based screens across more than a thousand of human cancer cell lines to identify genes required for tumor cell fitness. Within DepMap, gene essentiality is quantified using a dependency score (DS), in which increasingly negative values indicate a greater reduction in cell viability following gene knockout and therefore stronger cellular dependency.

Analysis of both the complete DepMap collection and liver cancer cell lines identified 26 conserved genes with a dependency score (DS) below -1.0, the commonly used threshold for strong dependency in DepMap analyses (Table 1, Figure 2). These genes were retained for further investigation. Notably, a large number of these 26 candidate genes belong to the same mitotic regulatory network, supporting their functional role in supporting the proliferative states of cells. Furthermore, four of the highest-ranking candidates (i.e., with the lowest DS), are currently being evaluated in Phase II and III clinical trials for the treatment of human malignancies [RRM2 (DS -2.5; Phase3; NCT02466971, PLK1 (DS 2.28; Phase2; NTC06398587), KIF11 (DS -2.23; Pahse2; NCT02092922), CDK1 (DS -1.99; Pahse2; NTC02688907)], providing independent support for the biological relevance of our strategy.

**Figure 2.**
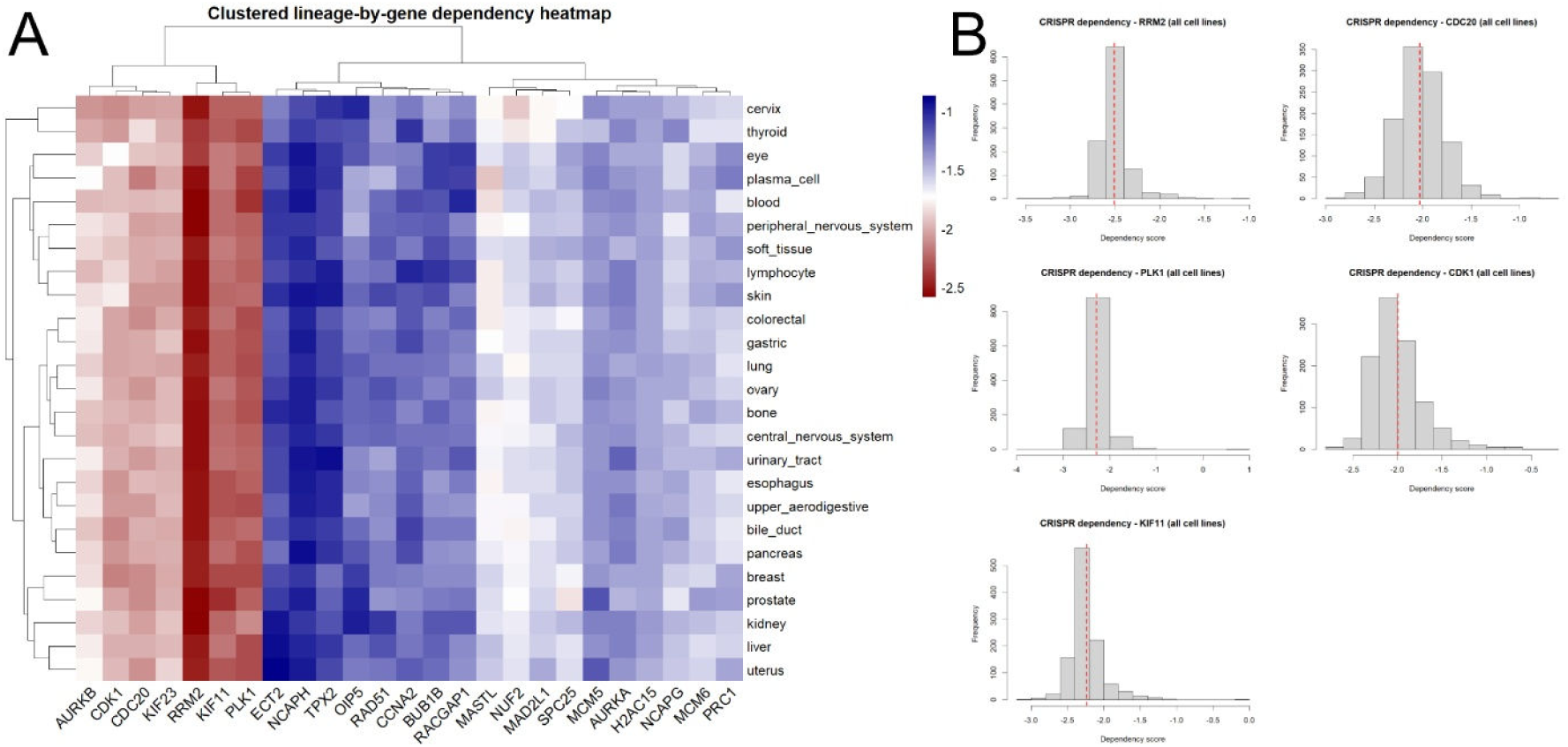
Functional dependency analysis of conserved candidate genes using DepMap. (**A**) eDepMap-generated hierarchical clustering of the 26 conserved candidate genes exhibiting strong dependency scores (DS < −1.0) across cancer cell lines representing 22 tissues, identified from the initial set of 380 queried genes. (**B**) Distribution of dependency scores for the five highest-ranking candidate genes across all DepMap cancer cell lines (>1,000 cell lines), illustrating their broad functional essentiality.

**Table 1.** Conserved tumor dependency genes prioritized through integrative computational analysis. List of the 26 candidate genes identified through sequential integration of differential expression analysis, functional dependency screening using DepMap.

| Gene | Dependency Score |
| --- | --- |
| RRM2 | -2.508837748 |
| PLK1 | -2.286522255 |
| KIF11 | -2.238001162 |
| CDC20 | -2.030719823 |
| CDK1 | -1.994233473 |
| KIF23 | -1.985713514 |
| AURKB | -1.866342169 |
| MASTL | -1.691319173 |
| NUF2 | -1.672701062 |
| MAD2L1 | -1.576458954 |
| SPC25 | -1.570621844 |
| PRC1 | -1.545050537 |
| NCAPG | -1.51256188 |
| MCM6 | -1.508903668 |
| H2AC15 | -1.429113715 |
| AURKA | -1.357555693 |
| MCM5 | -1.356593825 |
| RACGAP1 | -1.270485469 |
| OIP5 | -1.259324557 |
| RAD51 | -1.259062318 |
| BUB1B | -1.251855819 |
| CCNA2 | -1.211870948 |
| ECT2 | -1.112639325 |
| TPX2 | -1.087579811 |
| NCAPH | -1.000897626 |

### Conserved tumor dependency genes are transiently activated during regenerative liver responses

To investigate the biological context underlying the identified candidates, we examined whether these genes are associated with conserved proliferative programs activated during liver regeneration. Due to space limitations and to improve figure readability, the five highest-ranked candidates based on dependency scores (Table 1) were selected for graphical representation. We hypothesized that conserved tumor dependency genes identified across distinct HCC models are components of proliferative programs and should therefore be transiently induced during liver regeneration, returning to baseline expression once tissue homeostasis is restored. To test this hypothesis, we developed a dedicated version of our easyGEOarray (eGA) pipeline, termed eGA-light, which enables targeted analysis of user-defined gene sets across selected GEO datasets and generates integrated expression summaries.

The 26 candidate genes were analyzed across independent mouse models of liver regeneration and acute liver injury, including partial hepatectomy (PHx, GSE167034^12^), acute acetaminophen (APAP)-induced liver injury (GSE167032^12^), acute carbon tetrachloride (CCl₄)-induced liver injury (GSE222576^13^), and bile duct ligation (BDL, GSE166867^12^) (Figure 3). Across all models, candidate genes displayed a consistent temporal expression pattern, with basal expression in maintained at earlier time points, induction during the regenerative phase (i.e., day2-day6), and progressive return toward baseline levels upon restoration of tissue homeostasis.

**Figure 3.**
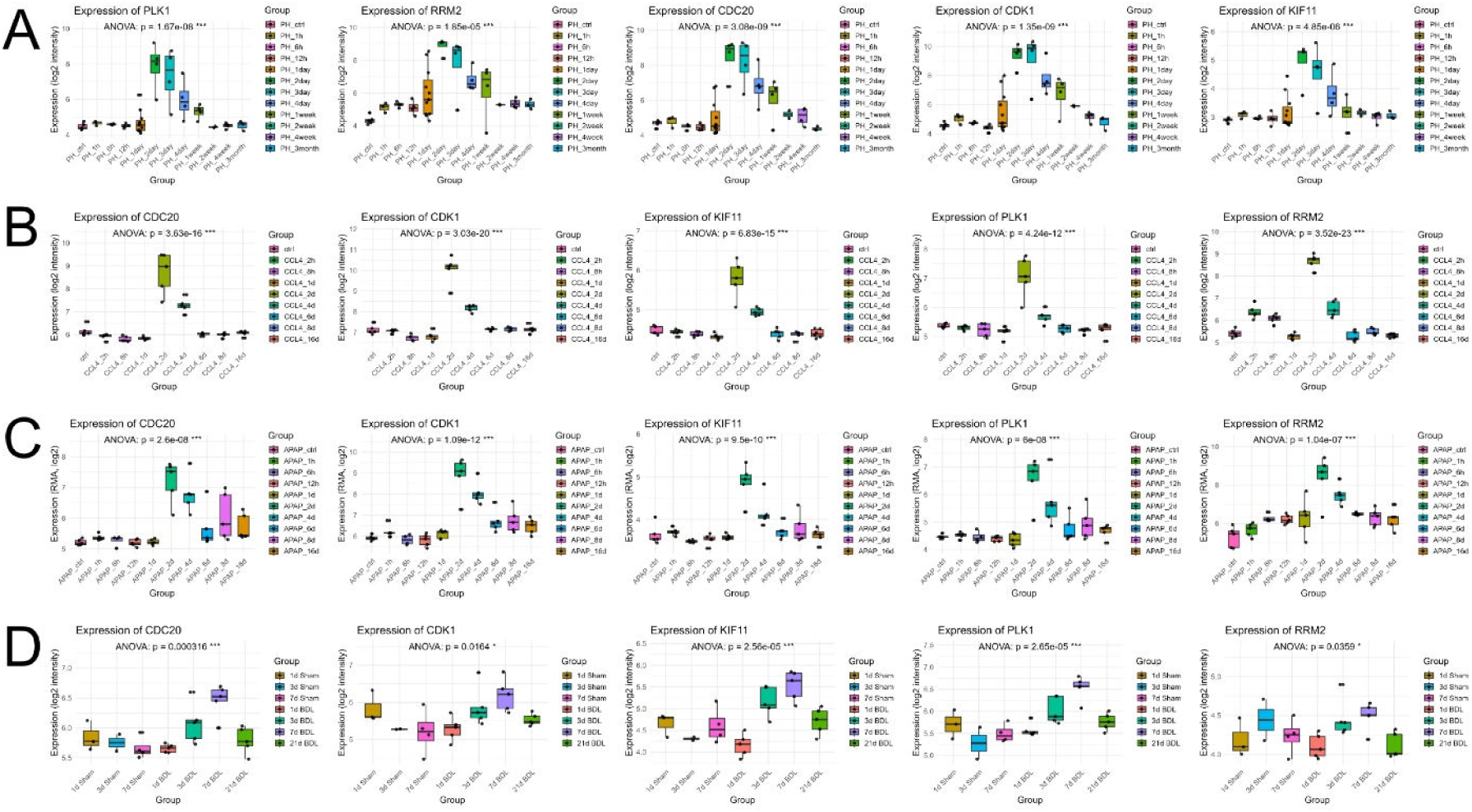
Selected candidate are transiently upregulated during liver regeneration. Expression profiles of the five highest-ranked candidate genes during experimental liver regeneration analyzed using the easyGEOarray-light (eGA-light) pipeline. Gene expression was evaluated in publicly available transcriptomic datasets following (**A**) partial hepatectomy (PHx; GSE167034), (**B**) acute carbon tetrachloride (CCl₄)-induced liver injury (GSE167033), (**C**) acute acetaminophen-induced liver injury (APAP; GSE167032), and (**D**) bile duct ligation (BDL; GSE166867). Candidate genes exhibit low basal expression in quiescent liver, significant transient induction during the regenerative phase (day 2 to day 6), and return toward baseline following restoration of liver homeostasis.

Together, these findings indicate that the candidate genes are components of conserved regenerative programs that are normally transiently activated to support liver repair but become persistently expressed in HCC, potentially contributing to tumor cell fitness through sustained proliferative signaling.

### Candidate genes remain persistently activated during chronic liver disease

To determine whether the transient activation pattern observed during liver regeneration is altered in chronic liver disease, the 26 candidate genes were further analyzed using eGA-light across independent models of chronic liver injury. Gene expression was evaluated in a chronic carbon tetrachloride (CCl₄) model analyzed at a single late time point (8 weeks, GSE141821^14^), a longitudinal CCl₄-induced fibrosis model including early and advanced stages of disease progression (2 and 12 months, GSE167216^12, 15^), and a mouse model of acute-on-chronic liver failure (ACLF, GSE298435^16^) (Figure 4). In contrast to the transient induction observed during regenerative responses, the candidate genes displayed sustained upregulation throughout chronic liver injury and ACLF. This persistent expression pattern suggests that proliferative programs normally switched off following tissue repair remain active during chronic liver disease.

**Figure 4.**
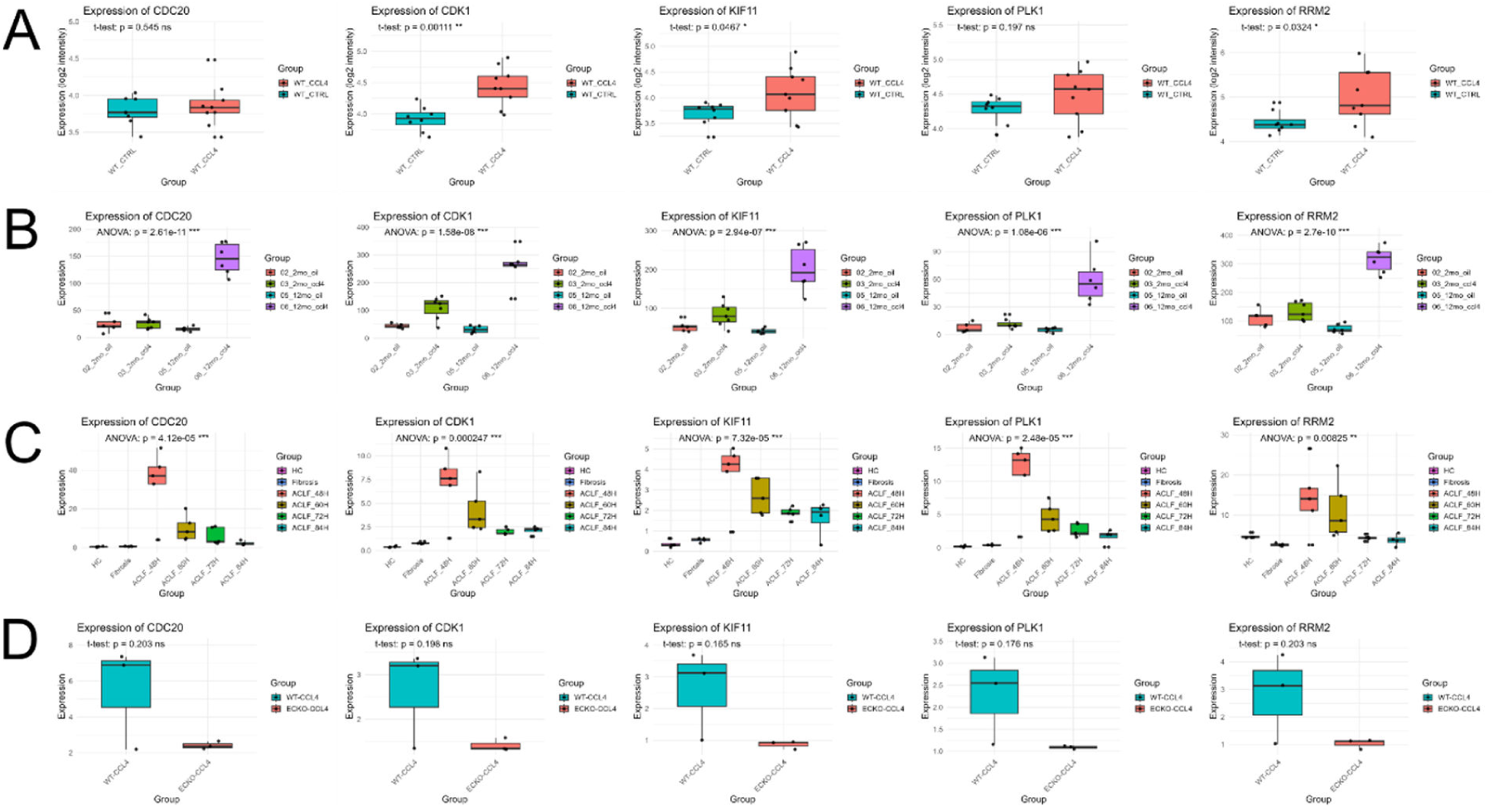
Sustained activation of candidate genes expression during chronic liver injury. Expression of candidate genes was analyzed using eGA-light in independent mouse models of chronic liver inhury, including (**A**) chronic carbon tetrachloride (CCl₄)-induced fibrosis (8 weeks; GSE141821), (**B**) longitudinal CCl₄-induced fibrosis (2 and 12 months; GSE167216), and (**C**) acute-on-chronic liver failure (ACLF; GSE298435). Unlike the transient expression observed during liver regeneration, candidate genes remained persistently upregulated throughout chronic liver injury. (**D**) Expression of candidate genes is reduced in primary liver sinusoidal endothelial cells (LSECs; GSE237859) isolated from endothelial-specific Tgfbr2 knockout (Tgfbr2^ECKO^) compared to LSEC isolated from control (Tgfbr2^f/f^) mice after CCl₄-induced liver fibrosis.

To further examine the relationship between chronic fibrogenic signaling and candidate gene expression, we analyzed transcriptomic data from primary liver sinusoidal endothelial cells (LSECs) isolated from endothelial-specific Tgfbr2 knockout (Tgfbr2^ECKO^) and control (Tgfbr2^f/f^) mice following CCl₄-induced liver fibrosis (GSE237859). Endothelial deletion of Tgfbr2 has previously been shown to attenuate liver fibrosis in this model. Consistent with our hypothesis, expression of the candidate tumor dependency genes was reduced in Tgfbr2-deficient LSECs compared with control cells (Figure 4B), suggesting that profibrotic TGF-β signaling contributes to maintaining the expression of these genes during chronic liver injury.

Together, these findings support the hypothesis that the identified candidates are components of conserved proliferative programs that are transiently activated during liver regeneration but become “constitutively” expressed during chronic liver disease and injury. The reduction in their expression following attenuation of TGF-β signaling further supports their association with pathological liver remodeling and identifies them as candidate molecular mediators linking fibrosis and tumor-associated proliferative programs.

### Candidate genes display a conserved response in human patients

To evaluate the clinical relevance of the identified candidate genes, we developed easyTCGAanalysis (eTA), a custom R pipeline for automated analysis of transcriptomic and clinical data from the TCGA-LIHC cohort. All 26 candidate genes were significantly overexpressed in HCC compared with non-tumorous liver tissue, and elevated expression of each gene was associated with reduced overall survival (Figure 5A).

**Figure 5.**
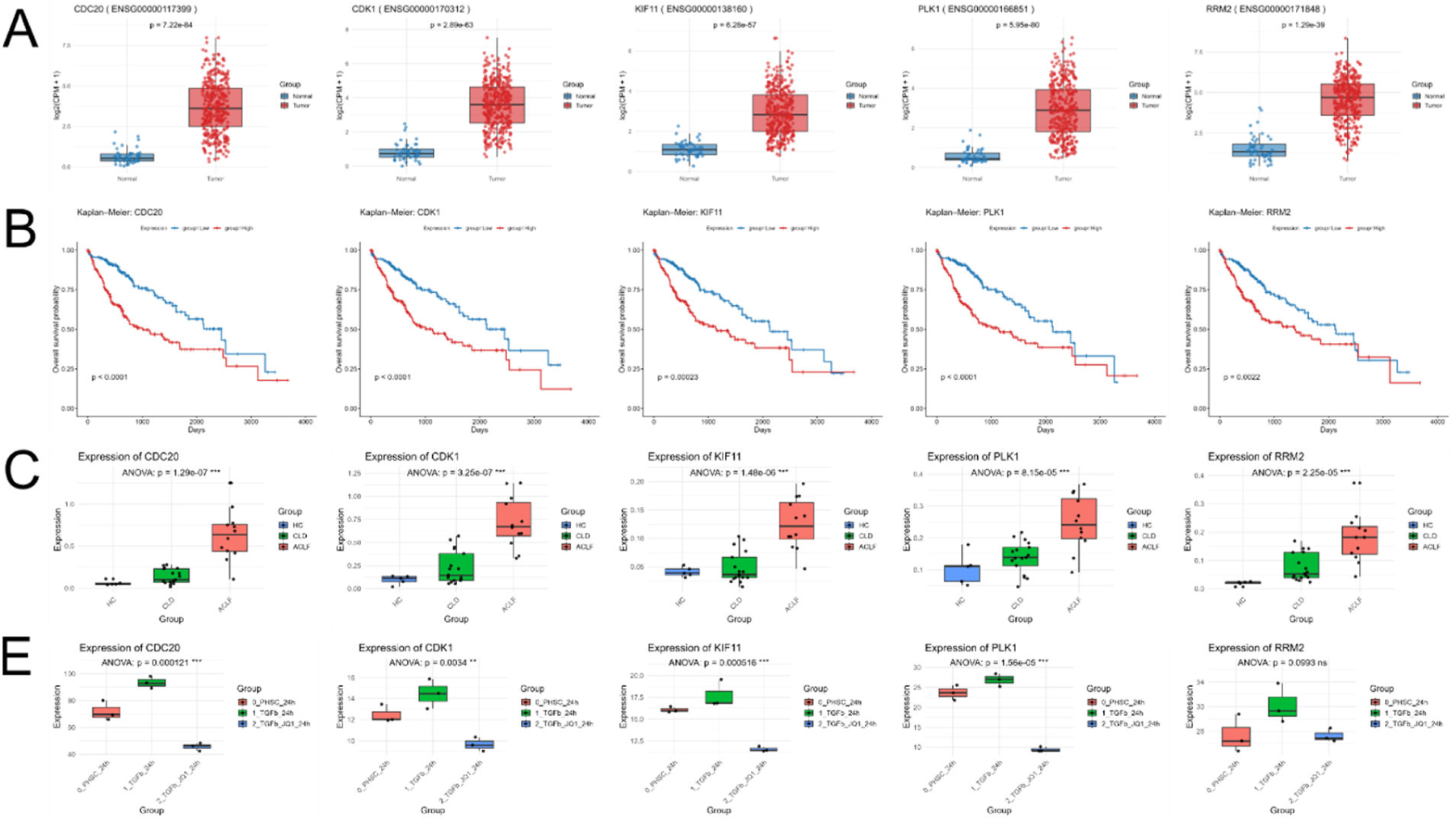
Validation of candidate genes in fibrotic patients with chronic liver disease. (**A**) Analysis of the TCGA-LIHC cohort using the easyTCGAanalysis (eTA) pipeline demonstrating significant upregulation of selected candidate genes in HCC (LIHC) compared with non-tumorous liver tissue. (**B**) Kaplan–Meier survival analysis showed that elevated expression of candidate genes was associated with reduced overall survival. (**C**) Expression of candidate genes in transcriptomic data from patients with cirrhosis and acute-on-chronic liver failure (ACLF; GSE298435). (**D**) Expression of candidate genes is significantly reduced in primary human HSC (PHSC; GSE117329) treated for 24h with TGFβ or TGFβ in combination with the BET bromodomain inhibitor JQ1.

We next investigated whether this conserved transcriptional program was also maintained during human chronic liver disease. Using eGA-light, transcriptomic data from patients with cirrhosis and acute-on-chronic liver failure (ACLF; GSE298435^16^) were analyzed. Consistent with the murine models, all candidate genes were significantly upregulated in both disease conditions (Figure 5B), demonstrating remarkable conservation of this proliferative program across species and throughout liver disease progression. To determine whether the TGF-β-dependent regulation observed in mouse was also conserved in human, we analyzed transcriptomic data from primary human hepatic stellate cells treated with TGF-β1 in the presence or absence of the BET bromodomain inhibitor JQ1 (GSE117329). JQ1, which suppresses BRD4-dependent TGF-β transcriptional programs and hepatic stellate cell activation^17^, significantly reduced the expression of multiple candidate genes compared with TGF-β1 treatment alone (Figure 5D), supporting a conserved TGF-β/BRD4-dependent regulatory mechanism in human hepatic stellate cells.

Finally, exploratory analyses of additional TCGA cohorts, including lungs (TCGA-LUAD), kidney (TCGA-KIRK), pancreatic ductal adenocarcinoma (TCGA-PAAD), and colorectal cancer (TCGA-COAD), revealed an overlapping trend toward increased expression of the candidate genes accompanied by poorer patient survival (Data not shown), suggesting that at least a subset of these dependencies may represent conserved vulnerabilities shared across multiple human malignancies.

## Discussion

The present study describes an integrative computational framework for the systematic identification and prioritization of conserved tumor dependency genes in hepatocellular carcinoma (Figure 6). By progressively integrating transcriptomic datasets from multiple murine models of liver cancer, functional dependency information from DepMap, experimental models of liver regeneration with chronic liver disease, and human transcriptomic and clinical datasets, we identified a conserved set of proliferation-associated genes that are consistently linked to liver disease progression and poor clinical outcome.

**Figure 6.**
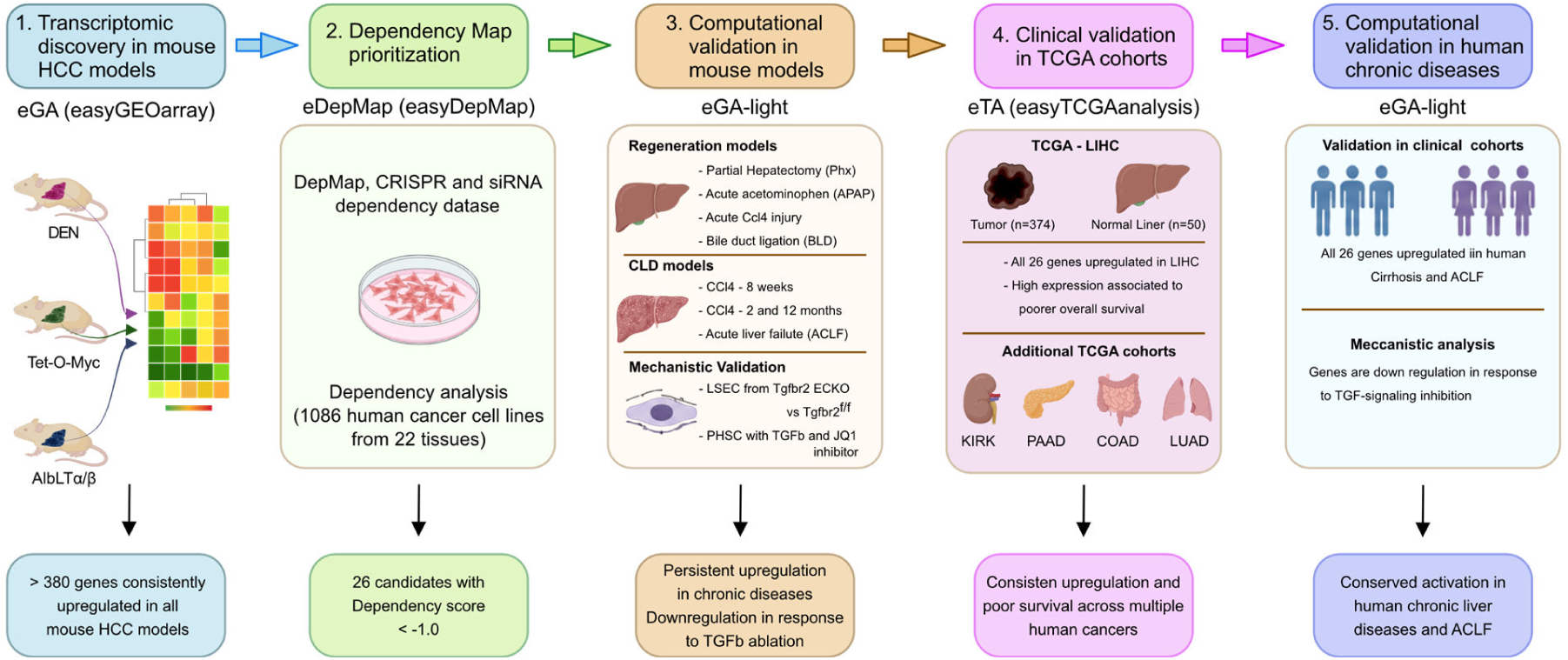
Computational workflow for the discovery of tumor dependency genes. Summary of the modular computational framework developed in this study. The workflow integrates transcriptomic analysis (eGA), functional prioritization (easyDepMap), biological (eGA-light), and clinical (eTA) validation to progressively identify conserved tumor dependency genes across murine models and human clinical data.

A central finding of this study is that the prioritized candidate genes exhibit a remarkably conserved expression pattern across diverse physiological and pathological contexts. During liver regeneration, these genes were transiently induced during the regenerative phase and subsequently returned to basal expression as tissue homeostasis was restored. In contrast, sustained expression was observed throughout chronic liver injury, acute-on-chronic liver failure, and HCC in both mice and humans. Together with their enrichment for cell cycle–related functions and strong dependency scores in human cancer cell lines, these observations suggest that the identified genes are components of conserved proliferative programs that are normally tightly regulated during liver repair but remain persistently activated during chronic liver disease and malignant transformation. This concept is summarized in the proposed biological model (Figure 7). Under physiological conditions (Figure 7A), regenerative programs are transiently activated following liver injury and are efficiently terminated once tissue repair is complete. During chronic liver disease and HCC (Figure 7B), these regulatory mechanisms appear to become uncoupled, resulting in persistent activation of proliferative pathways that support tumor cell fitness.

**Figure 7.**
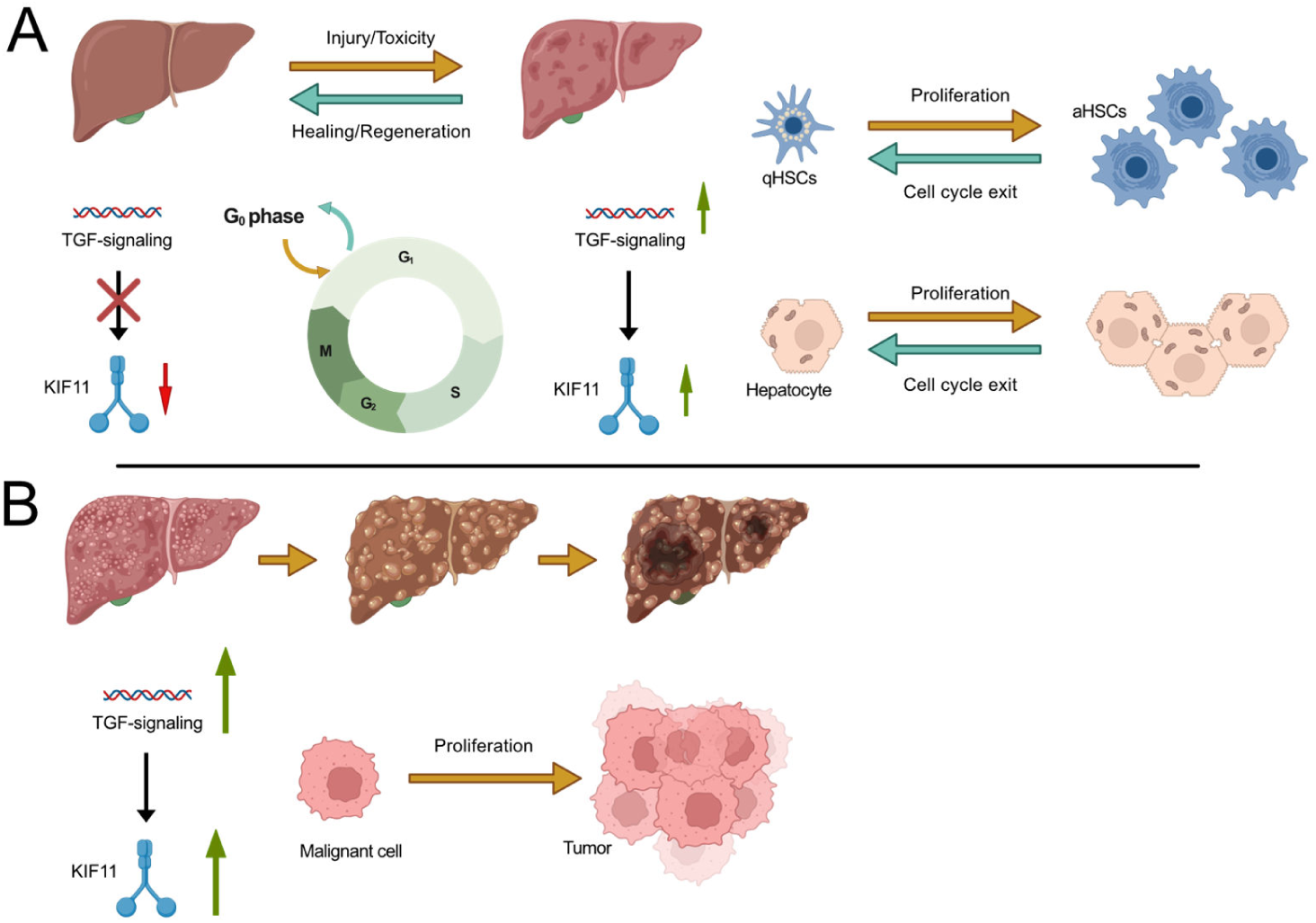
Proposed model illustrating the role of conserved tumor dependency genes during liver regeneration and hepatocarcinogenesis. (**A**) During acute liver injury, TGF–dependent regenerative signaling transiently induces proliferation-associated genes required for hepatocyte expansion, exemplified by KIF11. Once tissue homeostasis is restored, expression of these genes returns to basal levels. (**B**) During chronic liver injury and liver cancer, TGF–driven programs become persistently activated. Sustained expression of conserved tumor dependency genes supports continuous proliferative activity and tumor cell fitness, thereby creating potential therapeutic vulnerabilities that may be exploited through targeted inhibition.

These observations have important implications for therapeutic target discovery. Current targeted therapies for HCC largely focus on signaling pathways that are activated in specific molecular subgroups or etiological backgrounds. In contrast, the candidates identified in this study represent functional requirements that may be shared across genetically and etiologically distinct tumors. By integrating transcriptomic conservation with functional dependency screening, our framework prioritizes genes that are not only consistently expressed but are also predicted to be required for tumor cell survival. The findings that, several of the highest-ranked candidates genes identified in our analysis, (e.g., RMM2, PLK1, KIF11 and CDK1), are already being evaluated in phase 2 and 3 clinical trials for human malignancies, provides independent support for the biological relevance of the prioritization strategy. Furthermore, the observation that many candidate genes displayed similar expression and prognostic associations across multiple human cancer types suggests that at least a subset of these dependencies may represent Pan cancer vulnerabilities extending beyond primary liver cancer.

An important strength of the present study is the development of a modular computational workflow that integrates complementary public resources into a single analytical framework (Figure 6). Rather than relying on a single dataset or analytical platform, the workflow combines transcriptomic profiling, functional genomics, experimental disease models, and clinical validation through a sequential prioritization strategy. Each analytical step reduces the candidate space while increasing biological confidence, ultimately generating a manageable set of genes supported by multiple independent lines of evidence. Although developed for HCC, the individual modules are readily adaptable to other tissues and disease contexts, providing a flexible framework for the discovery of potential therapeutic targets in human malignancies.

Several limitations should be acknowledged. This study is based on the integration of publicly available datasets and therefore identifies biological associations rather than demonstrating direct causality. While functional dependency data strengthens candidate prioritization, experimental validation will be required to define the specific roles of individual genes in liver tumor development and therapeutic response. Furthermore, the upstream mechanisms driving the activation of these proliferative programs during chronic liver disease remain to be investigated.

In conclusion, the presented approach provides a resource for therapeutic target discovery and prioritization of novel molecular vulnerabilities for future validation and RNA-based intervention strategies.

## Material and Methods

### Computational analysis and software development

All computational analyses were performed using R-based workflows and publicly available datasets as described below. Data processing, normalization, statistical analyses, visualization, and integration of multi-omics datasets were performed using custom R scripts and established Bioconductor and CRAN packages. The complete list of datasets, accession numbers, software versions, and analysis parameters will be provided in the final peer-reviewed version of this manuscript.

To facilitate reproducibility and accessibility, the computational workflows generated in this study are being organized into a dedicated software suite, termed easyOmics, designed to provide a user-friendly framework for integrated omics data analysis and visualization. The fully documented release, including source code, user documentation, version information, and identifiers (repository link, release version, and DOI), will become available upon publication of the peer-reviewed manuscript. These resources are therefore not yet publicly accessible at the preprint stage.

## Figures were generated with Biorender

### Abbreviations

(HCC): Hepatocellular carcinoma,
(DepMap): Cancer Dependency Map,
(GEO): Gene Expression Omnibus,
(GO): gene ontology,
(TCGA): The Cancer Genome Atlas,
(DS): dependency score,
(RRM2): Ribonucleoside-diphosphate reductase subunit M2,
(PLK1): Polo-like kinase 1,
(KIF11): Kinesin-like protein KIF11,
(CDK1): Cyclin-dependent kinase 1,
(CDC20): Cell Division Cycle 20,
(CRAN): Comprehensive R Archive Network,
(LUAD): Lung Adenocarcinoma,
(LIHC): Liver Hepatocellular Carcinoma,
(PAAD): Pancreatic Adenocarcinoma,
(COAD): Colon Adenocarcinoma,
(TGF-β): Transforming Growth Factor Beta,
(TGFBR2): Transforming Growth Factor, Beta Receptor II,
(BRD4): Bromodomain-containing protein 4,
(ACLF): Acute-on-Chronic Liver Failure,
(LSEC): Liver Sinusoidal Endothelial Cells,
(CCl₄): carbon tetrachloride,
(APAP): acetaminophen,
(PHx): partial hepatectomy,
(DEN): Diethylnitrosamine,
(NTL): non-tumorous liver adjacent to tumors,
(CTRL): normal liver.

## Acknowledgment

The authors are grateful to the Department of Gastroenterology, Hepatology and Infectious Diseases at the University Hospital of Dusseldorf for providing a valuable test bed for the evaluation of the easyOmics computational framework.

